# Improved determination of protein turnover rate with heavy water labeling by mass isotopomer ratio selection

**DOI:** 10.1101/2024.06.04.597043

**Authors:** Jordan Currie, Dominic C. M. Ng, Boomathi Pandi, Alexander Black, Vyshnavi Manda, Jay Pavelka, Maggie P. Y. Lam, Edward Lau

## Abstract

The synthesis and degradation rates of proteins form an essential component of gene expression control. Heavy water labeling has been used in conjunction with mass spectrometry to measure protein turnover rates, but the optimal analytical approaches to derive turnover rates from the isotopomer patterns of deuterium labeled peptides continue to be a subject of research. Here we describe a method, which comprises a reverse lookup of numerically approximated peptide isotope envelopes, coupled to the selection of optimal isotopomer pairs based on peptide sequence, to calculate the molar fraction of new peptide synthesis in heavy water labeling mass spectrometry experiments. We validated this approach using an experimental calibration curve comprising mixtures of fully unlabeled and fully labeled proteomes. We then re-analyzed 17 proteome-wide turnover experiments from four mouse organs, and showed that the method increases the coverage of well-fitted peptides in protein turnover experiments by 25–82%. The method is implemented in the Riana software tool for protein turnover analysis, and may avail ongoing efforts to study the synthesis and degradation kinetics of proteins in animals on a proteome-wide scale.

**What’s new:** We describe a reverse lookup method to calculate the molar fraction of new synthesis from numerically approximated peptide isotopomer profiles in heavy water labeling mass spectrometry experiments. Using an experimental calibration curve comprising mixtures of fully unlabeled and fully labeled proteomes at various proportions, we show that this method provides a straightforward way to calculate the proportion of new proteins in a protein pool from arbitrarily chosen isotopomer ratios. We next analyzed which of the isotopomer pairs within the peptide isotope envelope yielded isotopomer time courses that fit most closely to kinetic models, and found that the identity of the isotopomer pair depends partially on the number of deuterium accessible labeling sites of the peptide. We next derived a strategy to automatically select the isotopomer pairs to calculate turnover rates based on peptide sequence, and showed that this increases the coverage of existing proteome-wide turnover experiments in multiple data sets of the mouse heart, liver, kidney, and skeletal muscle by up to 25–82%.

## Background

It is not possible to discern the relative contributions of synthesis and degradation to a change in protein concentration, and turnover (the combined processes of synthesis and degradation continue, for nearly all proteins, in the absence of any change in protein level - the ‘steady state’. Accordingly, the only way to assess turnover parameters is by monitoring flux of a label (whether stable or less commonly now, unstable, radioactive isotopes) through the protein pool. Stable isotope labeling can employ amino acids or simple metabolic precursors, such as heavy water or [^15^N] labeled ammonium ions.

Heavy water labeling, coupled with mass spectrometry, can be used to trace the synthesis and degradation kinetics of proteins in rodents and in humans ^1–13^. Under continued enrichment of heavy water D_2_O, deuterium (D or ^2^H) atoms are incorporated into non-essential amino acids during biosynthesis and metabolism. The deuterium labeled amino acids are in turn incorporated into nascent protein chains. The isotope incorporation rate over time reflects the rate of turnover of the protein pool. Water labeling is not readily compared to other forms of amino acid labeling, in which tracer amino acids are labeled consistently with a fixed number of ^13^C or ^15^N atom centers. These give a fixed mass offset per amino acid instance that is the same irrespective of the amino acid sequence, exemplified by SILAC or dynamic SILAC approaches. By contrast, heavy water labeling varies from one amino acid to another and thus, the degree of labeling of a peptide is sequence dependent. Moreover, whilst ^13^C or ^15^N amino acid labeling can be initiated with 100% of the amino acid pool being labeled, this is not feasible with water labeling-enrichments of 10% or less excess deuterium is usual. The combination of partial labeling and the variable number of labeled atom centers in each amino acid means that peptides exhibit complex labeling patterns showing a gradual trajectory of increasing and overlapping mass without the clear isotope separation typified by SILAC. The SILAC fixed mass offset greatly simplifies downstream data analysis, particularly as the labeled mass offset can mean that there is negligible contamination of the labeled isotopomer distribution with the unlabeled isotopomer distribution. With heavy water labeling, the isotopomer profile gradually shifts from unlabeled to labeled with considerable overlap of labeled and unlabeled profiles, leading to complex isotopic patterns (**Figure 1A**).

**Figure 1.**
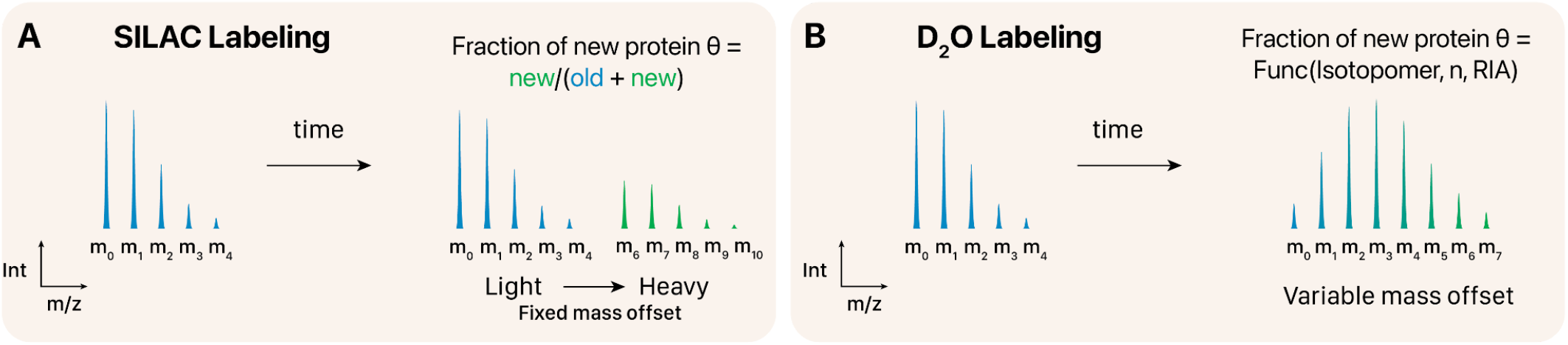
Data analysis in D_2_O labeling. Comparison of **A**. SILAC and **B**. heavy water labeling. SILAC labeling leads to fixed mass offset. The full replacement of the labeled residue with the heavy version with multiple heavy atom centers lends to relatively easy calculation of the molar fraction of newly synthesized proteins θ. Water labeling on the other hand is performed at low (< 20%) relative isotope enrichment and moreover leads to gradual shifts in the isotope envelope that are highly dependent on peptide sequences.

A critical step of the data analysis is to calculate the molar fraction of newly synthesized proteins, or fractional synthesis 0 ≤ θ ≤ 1, from the peptide isotope envelope of an MS1 spectrum. Commonly, this measurement is performed by considering the ratio of the mono-isotopomer (i.e., the first isotopomer, m_0_) over the complete isotope profile of the peptide m_0_/∑m_i_ The complete profile is typically computed from the intensities of the first six isotopomers (m_0_:m_5_) (**Figure 1B**). However, it should be noted that for longer peptides and depending on the enrichment level, m_6_ and higher isotopomer peaks can also be expected to be present at appreciable proportions. For instance, a peptide with 26 averagine residues will have 1% relative abundance in the m_6_ isotopomer prior to labeling, which rises to 10.1% assuming a conservative one label-accessible hydrogen atom per residue at 4.6% background deuterium enrichment.

Because of the gradual shift in isotopomer profile, the ratio of any pair of mass isotopomers contains information about the degree of deuterium enrichment. Partial mass isotopomer profiles (as low as a pair of isotopomers) could, in principle, be sufficient for calculating the fraction of newly synthesized proteins. Recently, Sadygov and colleagues derived closed-form analytical formulae for calculating the abundance ratios of the m_1_ to m_5_ mass isotopomers as a function of deuterium enrichment and amino acid labeling sites. These formulae permitted calculation of the most probable deuterium enrichment from the corresponding isotopomer ratio, from which the relative abundance of the monoisotopomer and turnover rates is calculated ^14^. The use of a subset of isotopomer ratios between two mass isotopomers (e.g., m_0_/m_1_) gave increased performance over using the cluster, as measured by the number of peptides whose isotopomer time series can be fitted well to a kinetic model (with R^2^ ≥ 0.9). The authors suggest that this is due to reduced chance of interfering isobaric contaminants in the isotopic cluster when only two isotopomer peaks need to be quantified. However, a limitation of this approach is that each isotopomer ratio calculation requires a separate combinatorics calculation, and not all formulae (m_6_ and beyond) are demonstrated.

Alternatively, the complete isotopomer profile of a peptide can also be calculated numerically with an isotope fine structure calculation algorithm such as IsoSpec2 ^15^ and enviPat ^16^, which resolves the isotopologues of compounds given their chemical compositions. The isotope envelope of a labeled peptide can be simulated by using the isotope fine structure calculators with a custom elementary composition table, which can resolve the probability (i.e., proportional abundance) of mass isotopomers beyond m_5_. Here, we show that the calculated isotopologue probability method can be applicable to heavy water based protein turnover analysis, and allows any arbitrary combination of isotopomer profiles to be directly queried numerically (e.g., m_1_/m_3_, m_1_/(m_0_+m_1_+m_2_), m_0_/Σ(m_0_:m_8_), etc.) without requiring the closed-form analytical solutions. A numerical lookup method to compare empirical isotopomer pair values to mixture spectra should then find the fractional synthesis of a peptide (**Figure 1C**), but to our knowledge, this approach has not been explored for heavy water labeling.

## Methods

### Cell Culture and Mass Spectrometry

For calibration samples, human AC16 cells (Millipore) were cultured in DMEM/F12 supplemented with 10% FBS and either 6% D_2_O (heavy labeled population) or 6% H_2_O (control population) at 37°C, 5% CO_2_. The cells were maintained in this medium for 3 passages, each passage with a split ratio of 1:8. This growth was estimated to constitute approximately 9 doublings of the cell populations. The cells were harvested by trypsinization, pelleted, washed once with phosphate buffered saline, and pelleted again before snap freezing in liquid nitrogen and storing at –80°C. At the time of processing each pellet was resuspended in 1 mL of RIPA buffer (Thermo Scientific) supplemented with Halt Protease and Phosphatase Inhibitor Cocktail (Thermo Scientific). Proteins were extracted with sonication in a Biorupter Pico (Diagenode) with settings 10x 30 sec on 30 sec off at 4°C. Insoluble debris was pelleted and removed from all samples by centrifugation at 14,000 **g**, 5 minutes.

Protein concentration of all samples was measured with Rapid Gold BCA (Pierce). Cell lysates from the D_2_O and H_2_O media populations were then combined in a labeling series expressed as the proportion of protein that was labeled with heavy water: 0, 0.125, 0.25, 0.375, 0.5, 0.625, 0.75, 0.875 and 1. The samples were trypsin digested using a modified version of the filter-aided sample preparation approach as previously described ^17^. A total of 50 µg protein per sample in 250 µL 8M urea was loaded onto Pierce Protein Concentrators PES, 10K MWCO (Thermo Scientific) pre-washed with 100 mM ammonium bicarbonate (Ambic). The samples were again washed with 8 M urea to denature proteins and remove SDS. The samples were washed with 300 uL 100 mM Ambic twice. The samples were then reduced and alkylated with final concentrations 5 mM dithiothreitol (DTT) and 18 mM iodoacetamide (IAA) for 30 minutes at 37°C in the dark. DTT and IAA were removed with centrifugation and the samples were washed 3× with 100 mM Ambic. Samples were digested atop the filters overnight at 37°C with mass spectrometry grade trypsin (Promega) at a ratio of 1:50 enzyme:protein. The following day samples were cleaned with Pierce C18 spin columns (Thermo Scientific) according to the manufacturer’s protocol. Eluted peptides were dried under vacuum and redissolved resuspended in 0.1% (v/v) formic acid.

The samples were analyzed on a Thermo Q-Exactive HF quadrupole-Orbitrap mass spectrometer coupled to a nanoflow Easy-nLC UPLC with the Thermo EasySpray electrospray ionization source. Peptides were separated with a PepMap RSLC C18 column 75 μm × 15 cm, 3 μm particle size (Thermo Scientific) with a 90 minute gradient from 0 to 100% pH 2 solvent B (0.1% formic acid in 80% v/v LC-MS grade acetonitrile). The mass spectrometer was operated in data-dependent acquisition mode with scans between m/z 200 and 1650 acquired at a mass resolution of 60,000. The maximum injection time was 20 ms, and the automatic gain control was set to 3e6. MS2 scans of the 15 most intense precursor ions with charge states of 2+ to 5+ were acquired with an isolation window of 2 m/z units, maximum injection time 110 ms, and automatic gain control of 2e5.

Fragmentation of the peptides was by stepped normalized collision-induced dissociation energy (NCE) of 25 to 27. Dynamic exclusion of m/z values was used with an exclusion time of 30 s.

### Fractional Synthesis Calculation

The determination of protein turnover rate through heavy water labeling requires interpretation of the pattern of isotope incorporation in heavy water labeled peptides to recover the fraction of newly synthesized protein as a function of the period of labeling. Isotope incorporation is commonly traced using the change in m_0_ (i.e., monoisotopomer)/∑m_i_ (i.e., the complete isotope profile) over labeling points. ∑m_i_ is typically only quantified for the first sixth isotopomers (i.e., i=0 to 5) ^2,8,18^. For the m_0_/∑m_i_ calculation (also referred to as m_0_/m_A_ for m “all” below), as the protein pool turns over, the isotopomer profile is assumed to traverse linearly from initial position, i.e., the theoretical natural distribution of experimentally unlabeled peptides, calculated from the biosphere abundance of isotopes of C, H, O, N, and S, toward the final asymptotic distribution that is determined by the relative isotope abundance and number of accessible labeling sites of the peptides. The initial coordinatecan be calculated given the absence (other than natural abundance) of heavy isotopes in any of the atoms in the peptide; the fully labeled asymptote, often theoretical, is modeled on the initial isotopomer profile, conditioned by complete incorporation of amino acids in which the deuterium abundance is as high as can be achieved given the water enrichment with deuterium.

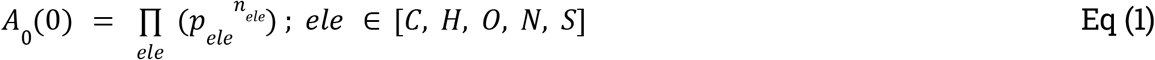

The plateau of A_0_ is calculated as the product of naturally occurring A_0_, times 1 – RIA to the number of deuterium accessible stable labeling sites n_l_, where RIA (relative isotope abundance) is the level of excess stable isotope labeling in the experiment.

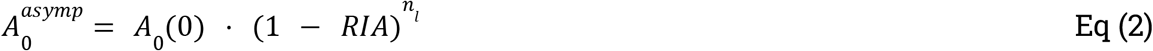

This could be refined by considering naturally occurring deuterium, but the background deuterium level is negligible and can be ignored. During labeling:

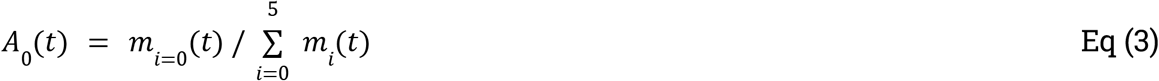

The fractional synthesis θis therefore calculated as:

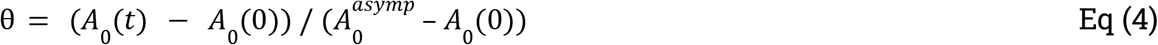

The intensity of the m_1_ isotopomer can be derived by closed-form equation as a function of excess deuterium enrichment. The intensity of the asymptotic abundance ratios up to the first sixth mass isotopomers (m_1_, m_2_, m_3_, m_4_, and m_5_) were reported by Sadygov and colleagues ^14^.

As an alternative to closed-form equations for peptide mass isotopomers, isotopic fine structure calculation algorithms can be used to resolve the isotopologues of any compound prior to and after deuterium labeling. Here we used IsoSpecR, which provides bindings to the IsoSpec2 ^15^ algorithm, to estimate the probability of isotopic combinations based on an elemental isotopic composition table. The heavier isotopes in peptides are largely driven by ^15^N and ^13^C; an isotope abundance of 0.003642 is used for ^15^N and 0.010788 is used for ^13^C in IsoSpecR. The ^13^C abundance is consistent with NIST values and corresponds to a δ^13^C of approximately –17 per VPDB standard (^13^C/^12^C of 0.011100), which is typical for biological materials. The isotopomer probabilities are then summed into mass isotopomers.

The pre-labeling and asymptotic predicted spectra are then summed into a series of composite spectra of any proportion of labeled and unlabeled peptide. These composite spectra are then compared to an experimental spectrum to estimate the fractional synthesis θfrom any isotopomer pairs of a peptide envelope:

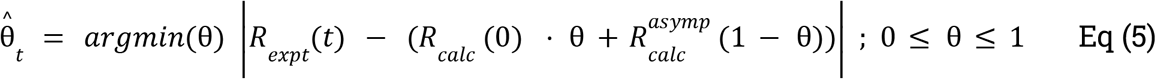

Where *R* is an applicable isotopomer ratio calculation, e.g., m_0_/m_1_, m_0_/Σ(m_0_:m_5_), etc.

### Kinetic Modeling and Statistical Analysis

Mass spectrometry data were searched against UniProt Swiss-Prot database ^19^ retrieved using Philosopher v.4.8.1 ^20^ on 2023-06-27 with added contaminants using Comet v.2022.01 ^21^ with typical parameters including: decoy_search = 1; peptide_mass_tolerance: 10.00 ppm; num_enzyme_termini = 1; isotope error: 0/1/2/3; fragment_bin_tol = 0.02; fragment_bin_offset = 0.0. Search results were post-processed using Percolator (crux-4.1 distribution) ^22^ with the following options: --decoy-prefix DECOY_; --overwrite T; --maxiter 15; --picked-protein. A peptide identification at FDR adjusted q value of 0.01 is considered a confident identification. Isotopomer intensity was extracted using Riana v.0.8.0 ^6^ to extract the intensity over time of the m_0_, m_1_, m_2_, m_3_, m_4_, and m_5_ peaks. Fractional synthesis was calculated as detailed above. The time series of fractional synthesis at different experimental time points was then fitted to a kinetic model to obtain the best-fit turnover rate constant (*k*_deg_) to explain the time series. For the re-analyzed adult mouse *in vivo* data, to perform kinetic modeling, we used the two-compartment model as described in Guan et al. ^23^ to find the best-fit *k*_deg_, with a high precursor equilibration rate constant (*k*_p_ = 3.0 d^−1^) that reflects the fast equilibration of heavy water with the protein precursor pool ^6,9^.

### Data and Code Availability

Riana is open-source and freely available on GitHub at http://github.com/ed-lau/riana. A visualization web app for isotopomer envelopment is available at http://heart.shinyapps.io/D2O_Isotope/. The re-analyzed data sets are available on ProteomeXchange at PXD029639 and PXD002870.

## Results

### Determination of fractional synthesis in heavy water labeling experiments using an isotopomer calculation algorithm

Using the isotopic fine structure calculation algorithm IsoSpec2, the asymptote isotope envelope of a peptide can be resolved by including an artificial element (H*) that represents the number of label accessible hydrogen atoms in the sequence with a probability of deuterium mass (2.0141 u) equal to the background enrichment level of heavy water in the experiment, which can be determined from direct measurement of body fluid by GC-MS or by direct fitting from peptide experimental data.^6^ A heavy water enrichment of 4.6% as from the mouse experiments in Hammond et al.^6^ is used in the analysis here. The number of H* atoms are then subtracted from the total number of hydrogen atoms. **Figure 2A** shows the simulated naturally occurring (pre-labeling) isotope envelope and asymptotic labeled envelope for three peptides: a relatively short peptide with few deuterium label sites (Y<u>F</u>D<u>L</u>G<u>L</u>PNR from mitochondrial isocitrate dehydrogenase, MW 1093.56, number of deuterium accessible labeling site n_l_ ∼ 13; essential amino acids underlined); a medium-length peptide with many non-essential amino acids and thus heavy water labeling sites (GQ<u>H</u>AAE<u>I</u>QP<u>L</u>AQS<u>H</u>AT<u>K</u> from myoglobin, MW 1785.91, n_l_ ∼ 47), and a very long peptide with very many deuterium labeling sites (<u>K</u>GS<u>IT</u>S<u>V</u>QA<u>I</u>Y<u>V</u>PADD<u>LT</u>DPAPA<u>TTF</u>A<u>HL</u>DA<u>TTVL</u>SR from ATP synthase subunit beta, MW: 3841.97, n_l_ ∼67). In the two longer peptides, particularly at asymptotic (4.6%) deuterium enrichment, there are appreciable m_6_ and higher probabilities present, which would indicate the typical m_0_/∑(m_0_:m_5_) calculation underestimates isotope incorporation (**Figure 2A**). This may be mitigated by expanding the conventional m_0_/∑m_i_ method to include for example integrating over up to the 10^th^ isotopomer (m_0_:m_9_) for peptides with more predicted labeling sites. However, integrating over multiple signal peaks leads to higher chances of contaminant isobaric peptides interfering with the quantification. The isotopomer envelopes of user-input peptide sequences can be visualized on a web app at http://heart.shinyapps.io/D2O_Isotope/.

**Figure 2:**
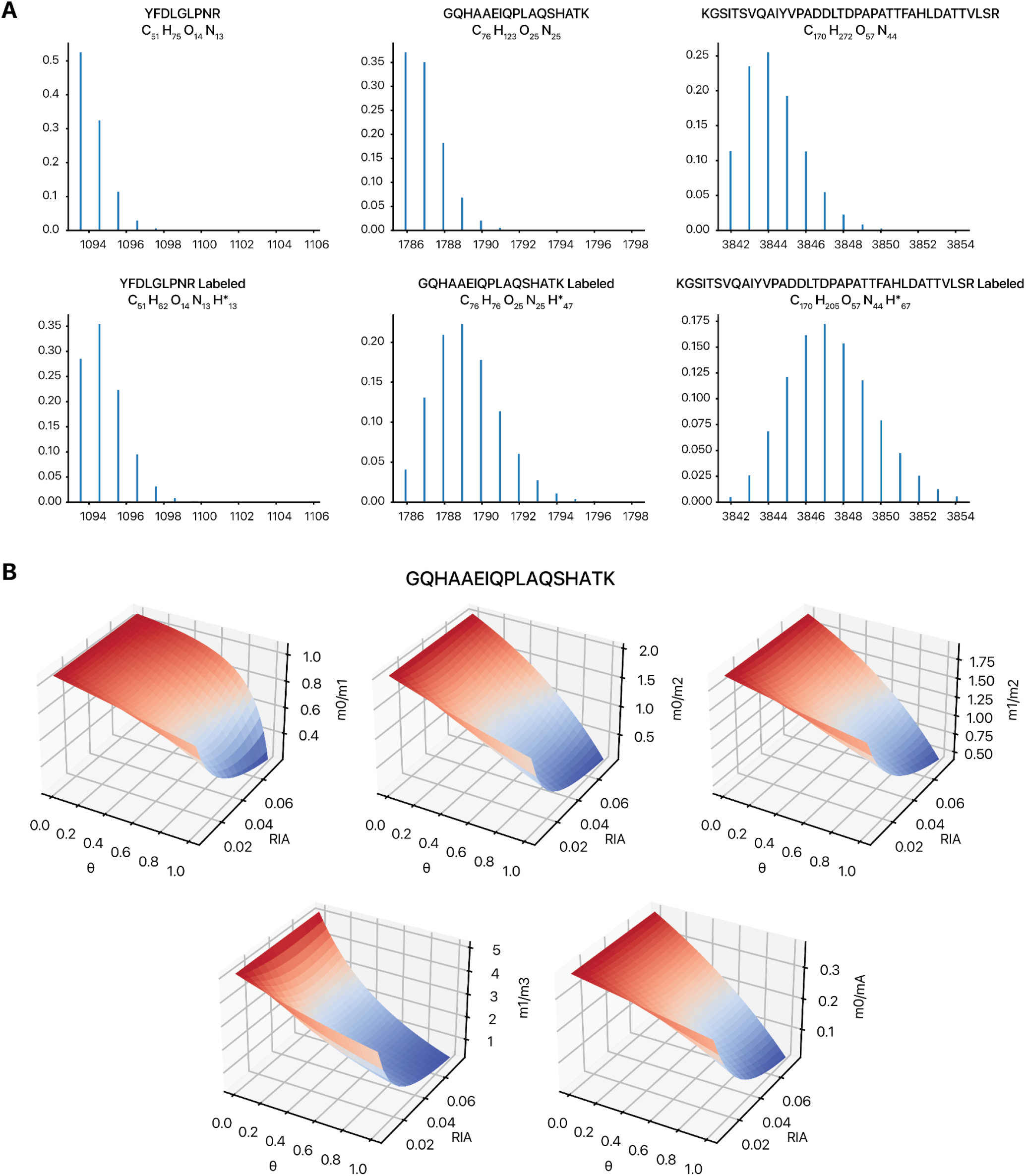
Simulated isotope envelopes in heavy water labeled samples. **A**. Simulated spectra showing the calculated isotopomer clusters based on element count and custom element count for three peptides that have been empirically quantified in a prior heavy water labeling data set (Hammond et al. 2022). (Left) A shorter peptide with few deuterium label sites, a medium length peptide with relatively high number of heavy water labeling sites, and a long peptide with many deuterium labeling sites. For each peptide, the top plot represents the simulated naturally occurring isotope envelope (pre-labeling), and the bottom plot represents the asymptotic isotope envelope following labeling with 4.6% relative isotope abundance (RIA) of heavy water. Appreciable m_6_ peaks can be seen in the long peptide prior to labeling, and medium and long peptides after labeling. **B**. Contour plots showing the relationship between fractional synthesis (θ), heavy water enrichment (RIA) and the isotopomer ratio of different measurements, including m_0_/m_1_, m_0_/m_2_, m_1_/m_2_, m_1_/m_3_, and m_0_/m_A_. Within each experimental RIA value under the shown range (x-axis), θ maps one-to-one to the shown ratios but with different spans.

The three dimensional contour plots for the same three peptides in **Figure 2A** are shown in **Figure 2B** and **Supplemental Figure S1**. The contours show the change of multiple isotopomer ratios as a function of fractional synthesis and the common range of heavy water labeling used in experiments (1 to 8%): m_0_/m_1_, m_0_/m_2_, m_1_/m_2_, m_1_/m_3_, and m_0_/m_A_ (i.e., Σ(m_0_:m_5_)), confirming that the ratios of each pair map to unique functional synthesis values given a particular known experimental precursor RIA. It can further be seen that the span and sensitivity of the isotopomer ratio vs. fractional synthesis relationship also depends on the background isotope enrichment (e.g., % deuterium in the system; RIA in the graphs below) and peptide sequence (number of labeling sites).

We then simulated the composite spectra by mixing linearly different proportions of the pre-label and post-label spectra. In this way, the empirically quantified ratios from any isotopomer pairs in an experiment (e.g., the intensity of the m_0_ peak divided by the m_1_ peak in the experiment) can be used to find the value of θ that would lead to the composite spectrum in Eq (5) that best matches empirical values. To evaluate how well the simulated composite spectra can be used to derive fractional synthesis from empirically measured isotope envelope, we re-processed the experimental data from a heavy water labeling experiments where proteins from the mouse heart were measured by mass spectrometry following up to ∼4.6% total body water deuterium enrichment in the animals at 12 time points for up to 31 days (**Figure 3B**).

**Figure 3:**
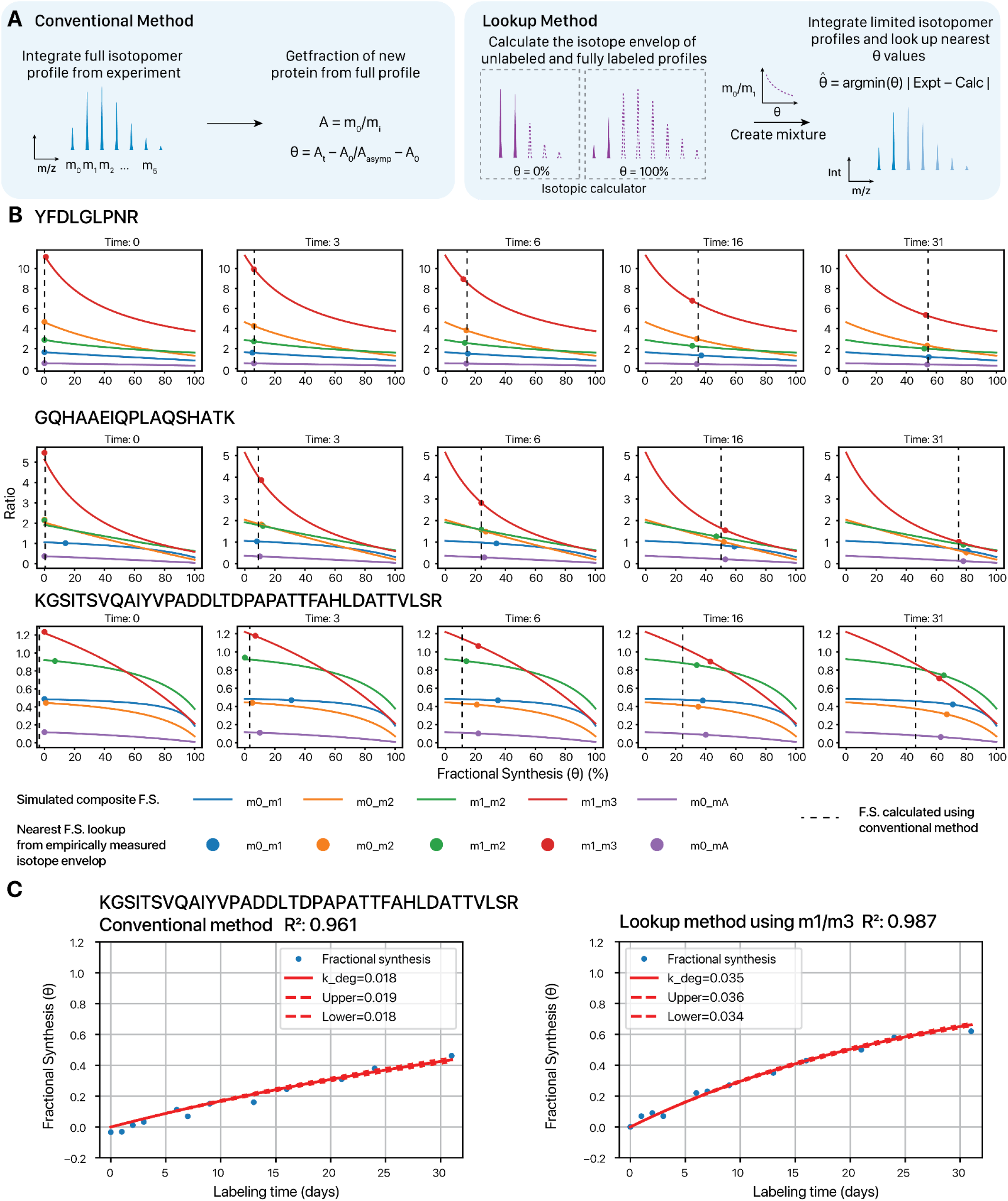
Estimating fractional synthesis from simulated composite spectra. **A**. (Left) The conventional method to calculate the molar fraction of new proteins from isotopomer profiles traces the fractional abundance of the first isotopomer (m_0_) without any heavy atom center over the entire isotope envelope. However, this approach requires integration over many isotopomers and is prone to isobaric contamination, or loss of m_0_ signal in long peptides. (Right) The lookup method uses a fine structure isotopic calculator to predict the isotope envelope for an unlabeled peptide (θ = 0) as well as a fully turned over labeled peptide (θ = 1). The virtual isotope envelopes are then mixed at different proportions to trace the relationship of any isotopic pairs with θ. This curve is then used to look up θ from the limited isotopic profiles (e.g., m_0_/m_2_ or m_0_/m_1_) from experimental data. **B**. Lines show the values of different isotopomer ratios (colors) in simulated composite spectra mixed from 0 to 100% of unlabeled and asymptotic peptide isotope envelopes. The data point shows the nearest estimated fractional synthesis from experimental data for the three peptides in consideration, from Hammond et al. 2022. The black dashed line shows the fractional synthesis value for the peptide at each time point as calculated using conventional methods in the original publication. **C**. Kinetic curve fitting of the peptide KGSITSVQAIYVPADDLTDPAPATTFAHLDATTVLSR with fractional synthesis values calculated from the conventional method (left) and the m_1_/m_3_ isotopomer ratio (right).

This led to several observations. First, in the three peptides above, it can be seen that the nearest θ method returns fractional synthesis rates that are consistent with one another and also consistent with the conventional method of fractional synthesis calculation used in the original study. This agreement appeared to be strongest for the short peptide with few labeling sites. In the two longer peptides with more labeling sites, the m_0_/m_1_ values deviated more from the other ratios, particularly at later time points of labeling where fractional synthesis is higher, the conventional method of “complete” isotopomer envelope starts to under-report true fractional synthesis compared to the partial isotopomer, which is consistent with appreciable m_6_ and above peaks in the isotope envelope. This is consistent with what is observed when the full (θ, t) series is used for curve fitting. The use of m_0_/m_2_ and m_1_/m_3_ improved the curve-fitting R^2^ value, and also derived a greater calculated turnover rate for the peptide over the conventional method (**Figure 3B**). Calculation of fractional synthesis from numerically resolved isotopomer profiles is sufficient for and may even improve kinetic modeling.

### Validation by a heavy water labeling calibration curve

To verify that the limited isotopomer ratios returned numerically accurate fractional synthesis values, we first set up a calibration curve experiment, where cultured human cells are cultured in 6% deuterium oxide for at least 9 doublings, estimated to lead to >99.8% complete labeling of all protein pools to their plateau deuterium relative isotope abundance ratios, i.e., all protein species are ∼100% labeled with 6% deuterium. This fully labeled pool is then mixed with non-deuterium labeled cell lysate at the fixed proportions of 0%, 12.5%, 25%, 37.5%, 50%, 62.5%, 75%, 87.5%, 100% to establish the ground truth of the simulation of fractional synthesis in each sample.

We focused on 1,672 peptides (from 666 protein groups) that were reliably identified in all nine mixture proportion experiments and were well behaved, as in having proportion of m_0_ steadily decreasing in a linear fashion with increasing proportion of labels, with a linear fitting R^2^ of 0.9, suggesting the isotopomer profiles for these peptides are accurately measured by the mass spectrometer and quantified by the software tool (**Figure 4A–B** m_0_/m_A_ curves). We then calculated the initial (unlabeled) and final (fully labeled) full isotopomer profiles using IsoSpec2, and numerically looked up the value of θ for the empirical isotopomer ratios using the mixed profiles as above, and compared the results with the ground truth mixture proportions. The resulting fractional synthesis curves largely maintained linearity and closely estimated the ground truth (two examples in **Figure 4A–B)** Overall, considering all 1,672 peptides, the θ lookup from m_0_/m_1_ showed a strong linear relationship close unity (y = 0.9688x + 0.041) and good agreement with the ground truth ratios (R^2^: 0.888) (**Figure 4C–D**) as does the m0/m2 ratio (y = 0.9847x + 0.024; R^2^: 0.890) (**Figure 4C, 4E**). We conclude that the isotopomer profile lookup method returned reliable fractional synthesis values.

**Figure 4:**
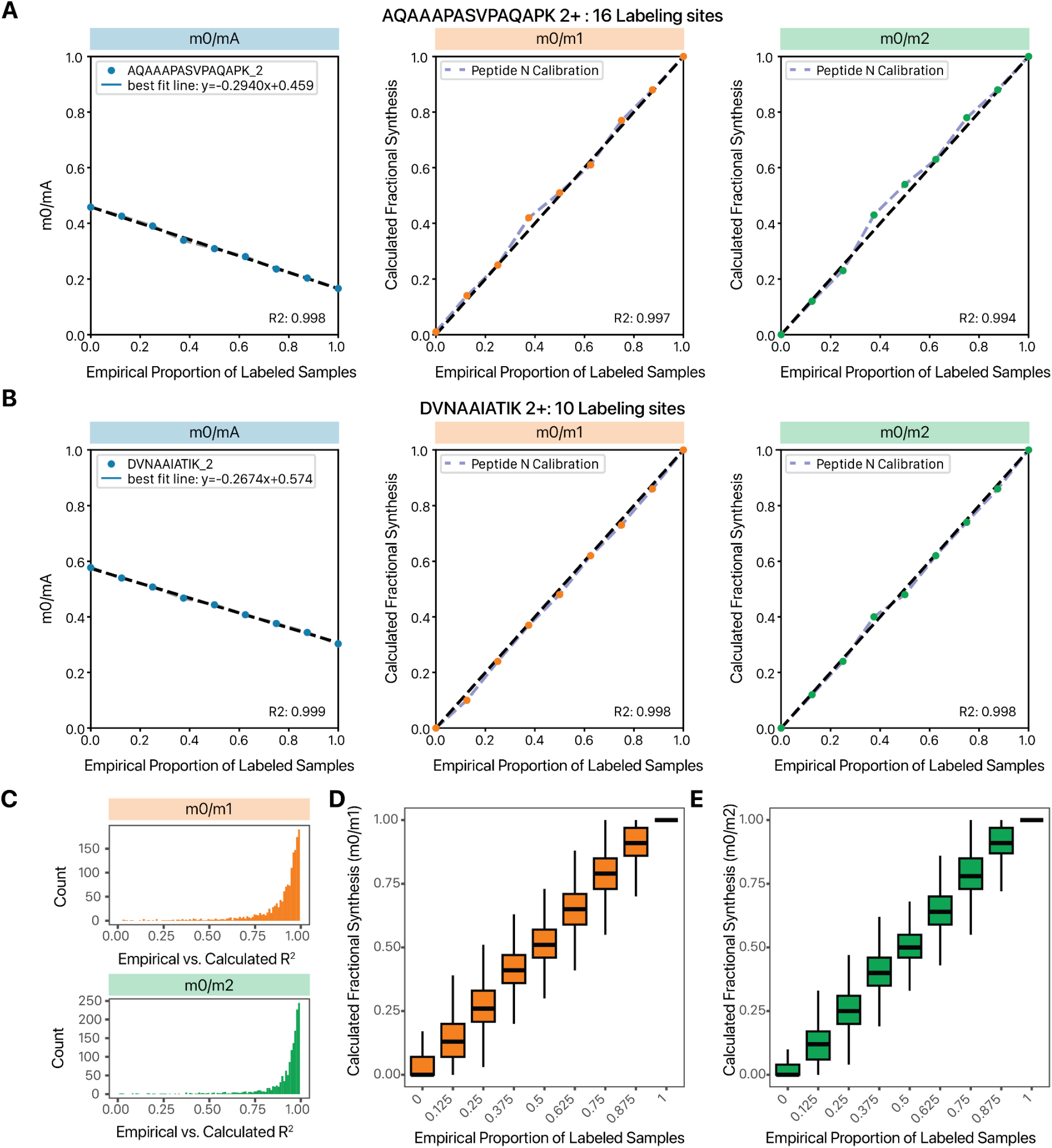
Isotopomer proportion calibration experiment using fully unlabeled and fully labeled samples. **A–B**. Human AC16 cells were labeled fully through 9 doublings in 6% D_2_O, then mixed with unlabeled cells at fixed proportion (0, 12.5%, 25%, 37.5%, 50%, 62.5%, 75%, 87.5%, 100% of the labeled cells). Two exemplary peptides (**A**. AQAAAPASVPAQAPK and **B**. DVNAAIATIK) are shown with a linear decrease in m_0_/m_A_ ratio from 0% to 100% labeling experiment (left). Using the isotopomer lookup method, we plotted the calculated fractional synthesis (y axis) from empirical m_0_/m_1_ (center) and m_0_/m_2_ (right) against the known mixture proportion (x-axis). **C**. Histogram of R^2^ values between empirical proportion of labeled samples vs. calculated fractional synthesis from the m_0_/m_1_ (top) and m_0_/m_2_ (bottom) ratios. Among 1,672 quantified peptide-charge pairs, 67.3% had R^2^ values ≥ 0.9 in the m_0_/m_1_ ratio and 74.5% had R^2^ values ≥ 0.9 in the m_0_/m_2_ ratio. **D-E**. Box plots showing the distribution of calculated fractional synthesis in **D**. m_0_/m_1_ and **E**. m_0_/m_2_ ratios. Center lines: median; boxes: interquartile range; whiskers: 1.5× interquartile range.

### Implementation of an automated isotopomer selection strategy to analyze protein turnover rates from four mouse organs

We next calculated the fractional synthesis profile toward the entire mouse heart data set for every distinct peptide-charge combination (hereafter peptides) that has been quantified in at least 6 labeling time points as in the original study. In total, we estimated the fractional synthesis and performed curve-fitting for 17,335 peptide time series. Using the conventional method, we derived the turnover rates of 6,851 distinct peptides that fit well to the kinetic model with R^2^ ≥ 0.9. We then compared this performance to calculations based on partial isotopomer profiles. Through discrete curve fitting for each of the m_0_/m_1_, m_1_/m_2_, and m_0_/m_2_ isotopomer ratio and selecting the best R^2^ value, we boosted the number of well-fitted peptides (R^2^ ≥ 0.9) by 40% (9,617 well fitted peptides), which is comparable to the gain reported by Deberneh et al. ^14^, who selected the isotopomer ratio pair to use by first performing curve-fitting on all quantified pairs, then choosing the one with the highest best-fit R^2^ value for each peptide. Although this approach leads to impressive gains, we reasoned that because the ratios derived from these isotopomer pairs would be expected to vary, the use of multiple θ series to elect for the best-fit is a case of multiple testing, where the R^2^ values are used first to compare the isotopomer pairs and select the best fit, then to report the final peptide goodness of fit. In addition, this post hoc selection does not allow a principled way to select the isotopomer ratio to use when only partial experimental data were collected or in experimental design where it is only possible to collect samples from one labeling time point. We therefore wonder whether the fitting could be improved using *a priori* rules, without relying on comparison of R^2^ values after curve fitting. Contrary to the report by Debeneh et al., the isotopomer ratios had unequal contributions to fitting improvements, with m_0_/m_2_ being the ratio that led to best R^2^ in 43.6% of the peptides, followed by m_1_/m_3_ (26.7%), then m_0_/m_1_ (21.4%) then m_1_/m_2_ (8.3%) (**Figure 5A**), which we attribute to differential sensitivity of the isotopomer pairs to θ in different peptides (**Figure 2**)

**Figure 5:**
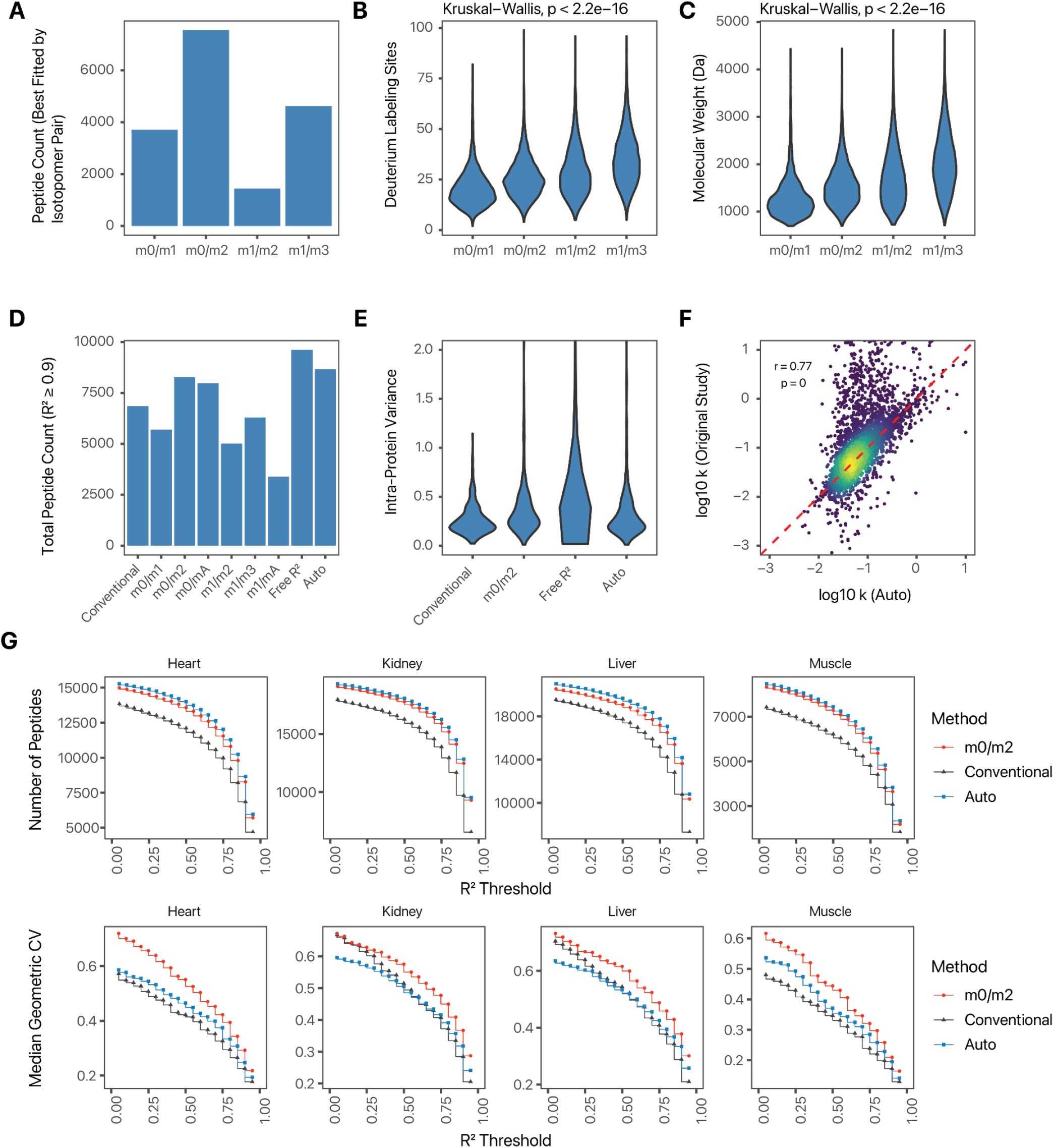
Comparison of number of quantifiable peptides at different R^2^ thresholds using different isotopomer pairs. **A**. Bar charts showing the number of peptides (y-axis) across four categories where a particular isotopomer ratio (x-axis) provides the best fit to the kinetic model over the other ratios. **B**. Boxplots showing the number of labeling sites in the peptides across each category. **C**. Boxoplots showing the molecular weight of peptides across each category. **D**. Total number of well-fitted peptides (R^2^ ≥ 0.9) in the Hammond et al. 2022 mouse heart data when using different fractional synthesis calculation methods (“Free R^2^” refers to picking the best R^2^ out of four isotopomer pairs after curve-fitting with each of them; “Auto” refers to rule-based prior selection, see text for details). **E**. Intra-protein variance, defined as the geometric coefficients of variance of *k*_deg_ of peptides mapping uniquely to the same protein, calculated from well-fitted peptides using different fractional synthesis calculation methods. **F**. Scatterplot showing the derived *k*_deg_ values in the “Auto” selection method over the values reported in the original study. **G**. Number of peptides (y) passing different R^2^ threshold (top) and the resulting intra-protein variance (bottom) at each threshold across four experiments (heart, kidney, liver, and muscle) from Hammond et al. 2022, when calculated using three different methods: conventional m_0_/m_A_ (gray), m_0_/m_2_ (red), and Auto (blue).

Notably, there is a strong relationship between the isotopomer ratios that lead to the best fit with the number of deuterium accessible labeling sites n_l_ (**Figure 5B**) and implicitly, the molecular weight (**Figure 5C**) of the peptides (Kruskal-Wallis P < 2.2e–16).

This may be because for short and medium peptides, the m_0_/m_2_ ratio spans the widest range of ratios across θ (**Figure 3A**) and high peak intensity for accurate integration, both of which improve fractional synthesis calculation. On the other hand, for very long peptides with many labeling sites, the m_0_ isotopomer intensity has a very low relative abundance that renders it less effective in accurate θ estimation (**Figure 2A**).

We therefore derived a simple isotopomer selection heuristics, where peptides with < 15 labeling sites are quantified using m_0_/m_1_, peptides with 15 ≤ labeling sites ≤ 35 are quantified with m_0_/m_2_, and peptides with >35 labeling sites are quantified with m_1_/m_3_ isotopomer ratios. In the mouse heart data set, this approach modestly outperforms quantifying with m_0_/m_2_ alone, and was able to improve on the conventional method by 27% (8,666 well fitted peptides belonging to 1,488 protein groups, vs. 6,851 peptides in 1,219 protein groups) without freely picking the best fit R^2^ values from multiple isotopomer pairs for each peptide (**Figure 5D**). The method increases the peptide multiplicity of proteins in the results, bringing the average number of quantified peptides per unique protein from 5.5 to 6.0. At the same time, this method was able to reduce the intra-protein variance over the m_0_/m_2_ or the best-R^2^ method (**Figure 5E**). Finally, the best-fit *k*_deg_ values were similar in numerical values to the conventional method derived *k*_deg_ values from the original study, so this method did not introduce an overall bias (**Figure 5F**).

We next applied the heuristic isotopomer selection method to data from three additional experiments, from two fast-turnover tissues (mouse liver and mouse kidney) and one slow-turnover tissue (mouse skeletal muscle). In each of the tested data sets, we observed a consistent increase in the number of well fitted peptides across multiple R^2^ thresholds (**Figure 5G**). The automatic isotopomer selection method led to 27% (8,666 vs. 6,851 peptides), 32% (12,830 vs. 9,707 peptides), 31% (14,190 vs 10,797 peptides), and 25% (3,835 vs 3,081 peptides) gain at R^2^ ≥ 0.9 for the heart, kidney, liver, and muscle data. The gain was greater for the two high-turnover tissues, but led to substantial increase in the depth of the turnover measurements in all four tissues than was published. In the liver for instance, the selection method led to more proteins (1,631 vs. 1,370) with well-fitted turnover rate information based on the same R^2^ ≥ 0.9 threshold. These proteins include those with liver biased expression and function, such as cytochrome P450 monooxygenases 2C37 (CYP2C37) 2C44 (CYP2C23), which are involved in polyunsaturated fatty acid metabolism, with measured turnover rates of 1.15 and 1.23 /d (R^2^ 0.961 and 0.903), respectively. Hence, the improved analysis can lead to more protein turnover information being recovered from existing data sets.

Finally, we applied the method to a separate data set of mouse hearts of the A/J and BALB/c strains, with or without isoproterenol-induced cardiac hypertrophy ^2,11^. This data set was generated on an Orbitrap Elite instrument in FT/IT mode, with substantial fractionation consisting of biological subcellular fractionations into the cytosolic, mitochondrial, and nuclear enriched fractions followed by two-dimensional peptide fractionation. The method of automated isotopomer selection followed by reverse lookup fractional synthesis calculation again led to substantial increase in the coverage of well-fitted (R^2^ ≥ 0.9; identified at ≥4 time points) peptides of 59%–82%, while keeping intra-protein variance under control.

**Table 1.**
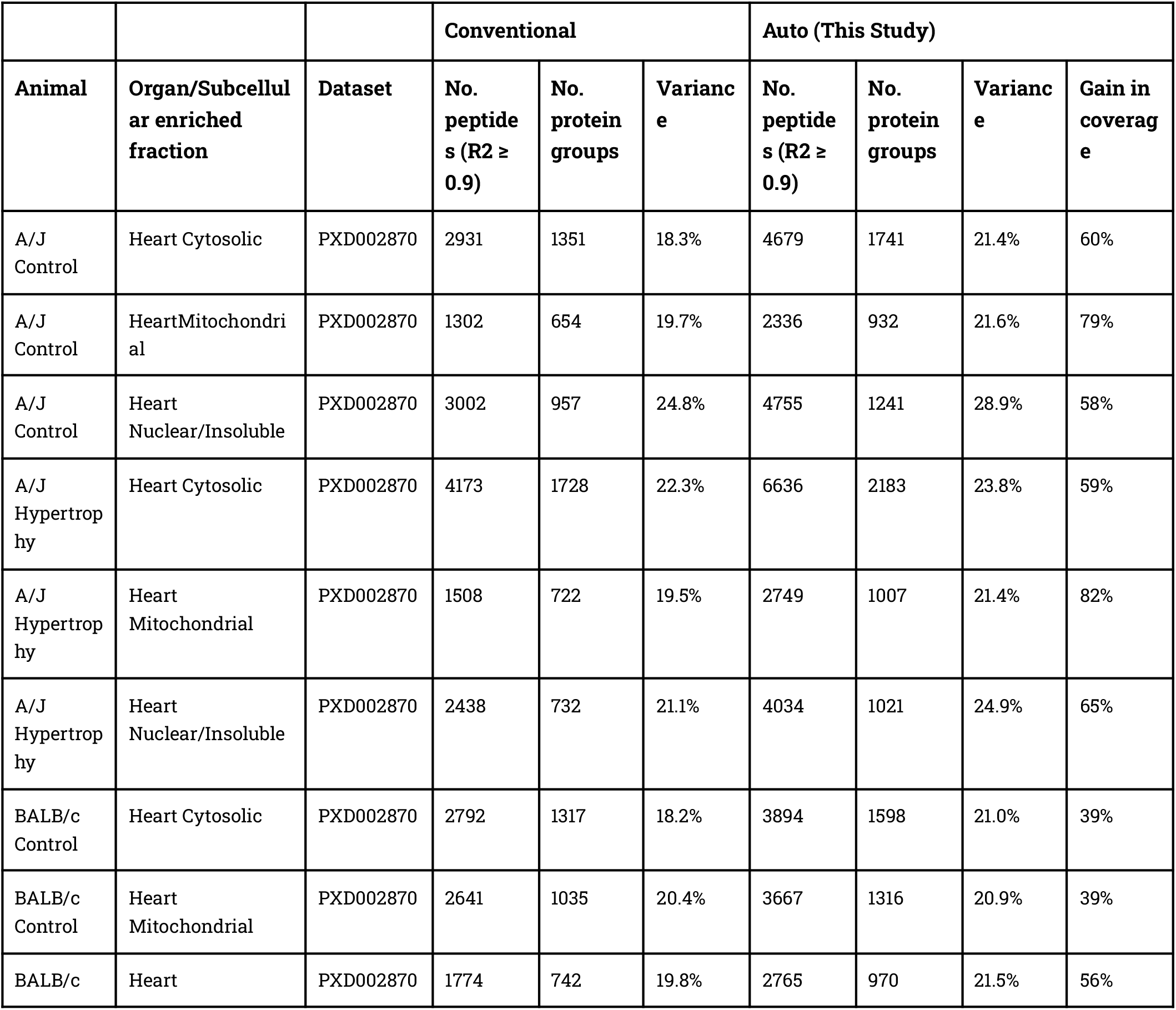
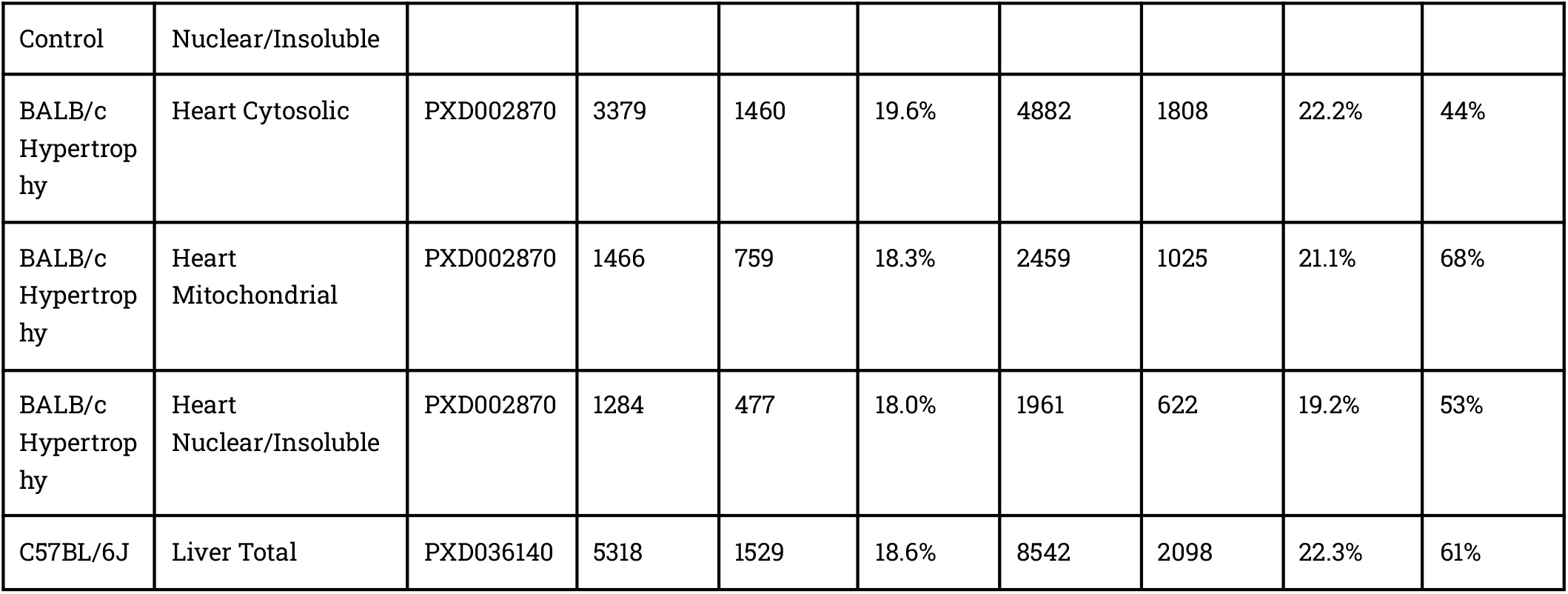
Increasing the depth of protein turnover coverage in 13 additional sets of turnover time series data from 3 mouse strains across 2 studies. Peptides quantified at more than half of the time points in each study were admitted for curve-fitting. Variance is calculated as the median of the geometric CV of proteins with 3 or more unique peptides fitted to the kinetic model at R^2^ ≥ 0.9.

## Discussion

Our group and others have applied heavy water labeling to examine the regulation of protein turnover in various adult animals and disease models ^1,4,8,11^. Compared to more commonly employed SILAC experiments, heavy water labeling has the advantages of quick precursor equilibration, low cost, and bio-orthogonality, and being applicable to many different types of tissues and animals ^5^. However, wider applications continue to be hurdled by the relative complexity in the interpretation of spectral data to extract the fraction of newly synthesized proteins. A heavy water labeled peptide can be considered as a polymer of multiple deuterium-accessible labeling sites and the isotopomer profile is a summation across the combinatorial possibilities of site labeling, factoring in number of labeling sites and the probability of incorporation of a deuterium atom (which is related, in turn to the precursor water enrichment). Mass isotopomer distribution analysis shows that the ratios of isotopomer pairs contain information regarding the relative isotope enrichment and molar fraction of synthesized peptides ^24^. Recent work, notably by Sadygov and colleagues, showed that “limited isotopomers” (using specific subsets of isotopically labeled ions rather than the full isotope envelope) can be applied to heavy water labeling ^18^. Here, we build on these prior works in three ways, by reporting: (1): a straightforward “reverse lookup” method to find the molar fraction of new synthesis from numerically approximated peptide isotopomer profiles in heavy water labeling studies; and (2): a rule-based isotopomer pair selection method that depends on total deuterium-accessible labeling sites in a peptide.

Numerical calculation of partial isotopomer profiles using isotopic fine structure algorithms allowed different isotopomer ratios from empirical spectra to be matched to the simulated composite spectra to find the fractional synthesis values of heavy water labeled peptides. Here, we provide corroborating evidence that the use of isotopomer pairs is sufficient for fractional synthesis calculation and improves upon the conventional “complete” isotopomer profile method. Moreover, we find that the m_0_/m_2_ ratio increases goodness-of-fit (R^2^) to the kinetic model in the greatest number of peptides, but does not perform as well for long peptides or peptides with great numbers of deuterium accessible labeling sites. We derived a simple heuristic isotopomer selection strategy where peptides with the number of accessible labeling sites n_l_ < 15 are quantified with m_0_/m_1_, 15 ≤ n_l_ ≤ 35 with m_0_/m_2_, and n_l_ > 35 peptides are quantified with m_1_/m_3_. We show that this simple selection strategy based on *a priori* defined rules is able to boost the number of peptides with well-fitted isotopomer trajectory over time to the kinetic model (R^2^ ≥ 0.9) substantially. Interestingly, we find that this approach of looking up the nearest simulated fractional synthesis in the calculated isotopologue profiles improves upon the conventional method of monoisotopomer relative abundance calculation, even when identical mass isotopomers (i.e., m_0_/Σ(m_0_:m_5_)) are used (**Figure 3D**). This suggests that other factors may also contribute to the improved turnover profiling coverage beyond the reduction of isobaric contaminant peptides.

These factors may include, firstly, the composite lookup method (Eq (5)) being bounded to 0 ≤ θ ≤ 1, whereas the conventional calculation (Eq (4)) is not and can return negative θ values unless additional boundary conditions are set. This can happen for instance when the proportion of the monoisotopomer in the complete envelope is lower than pre-labeling naturally occurring values or higher than the theoretical asymptotic probability due to measurement errors. Secondly, for longer peptides, the use of the first six isotopomers in the conventional calculation would under-estimate θ because the m_6_ to m_8_ peaks have appreciable intensity and hence m_0_/m_A_ loses linearity over θ. This is not the case in the lookup method, because the contributions of m_6_+ peaks are accounted for in both the empirical values and the composite simulated spectra. Thirdly, as noted above, different isotopomer ratios span different ranges and hence would offer differential sensitivity in quantification, whereas for longer peptides the m_0_ peaks are diminutive even prior to labeling and may not lend to accurate measurements. Taken together, we propose that a combination of factors likely contribute to the advantages of partial isotopomer profiles whether calculated from closed-form equations or from isotopic fine structure algorithms. Indeed, different isotopomers appear to best support goodness-of-fit to kinetic models for peptides with differential molecular weights or numbers of deuterium labeling sites. Overall, the described method of a priori selection of isotopomer pairs for fractions synthesis calculation based on peptide properties improved on conventional methods by as much as 82% when used to reanalyze 16 existing D_2_O labeling experiments from across data sets from different groups using different instrument types. This strategy is implemented in Riana, an open-source Python software for protein turnover quantification analysis compatible with heavy water and amino acid labeling.

## Acknowledgments

We thank Prof. Robert J Beynon at the University of Liverpool for the helpful discussion and editing of the manuscript. This work was supported in part by NIH award R35-GM146815 and the University of Colorado SOM Translational Scholar Research Program to E.L; and NIH awards R01-169473, R01-HL141278, and R01-GM144456 to M.L.

## Supplemental Information

Improved determination of protein turnover rate with heavy water labeling by mass isotopomer ratio selection

## Supplemental Figures

**Supplementry Figure S1.**
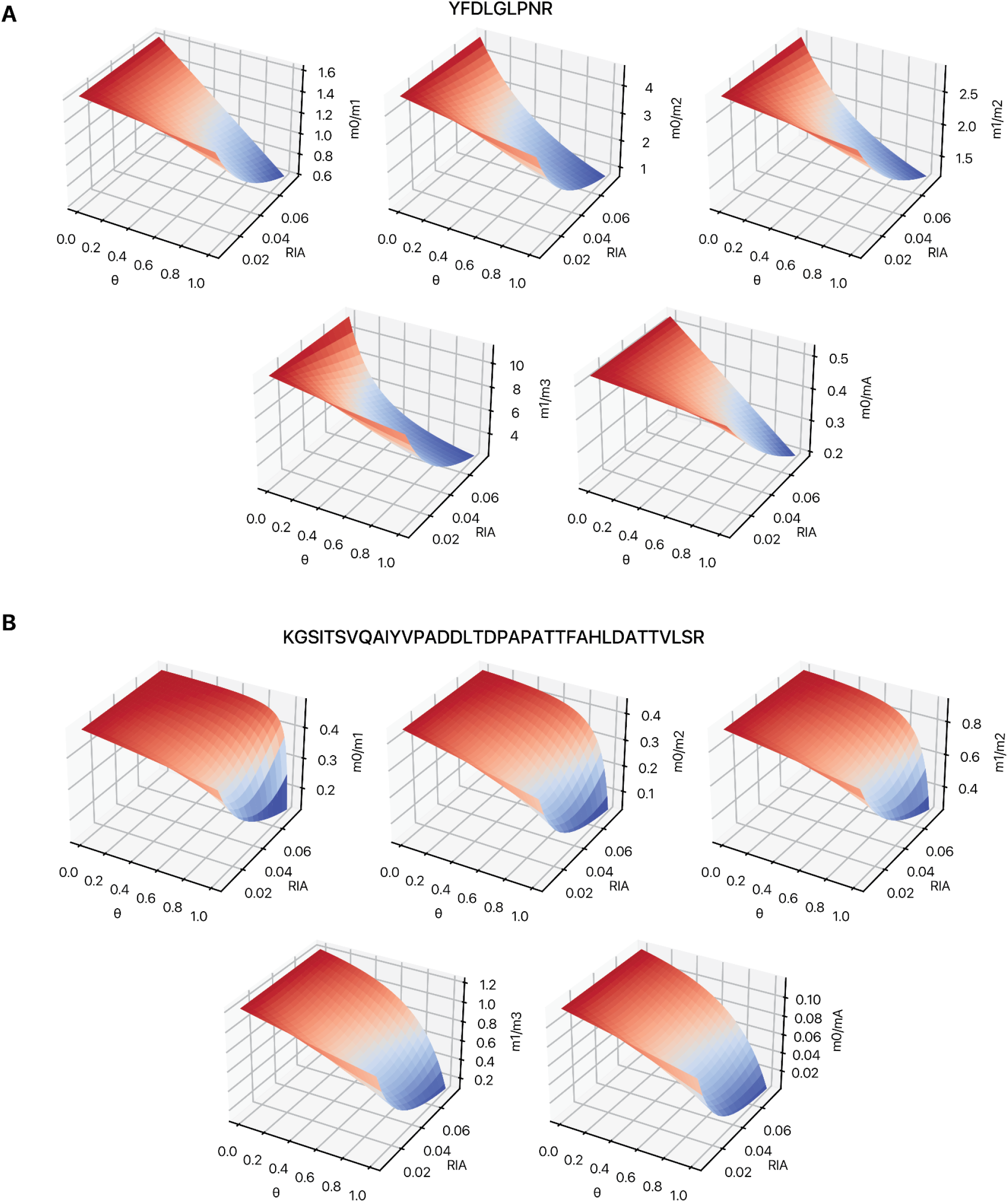
Additional simulated isotope envelopes in heavy water labeled samples. Contour plots showing the relationship between fractional synthesis (θ), heavy water enrichment (RIA) and the isotopomer ratio of different measurements, including m_0_/m_1_, m_0_/m_2_, m_1_/m_2_, m_1_/m_3_, and m_0_/m_A_, for two additional peptides corresponding to Figure 2: YFDLGLPNR and KGSITSVQAIYVPADDLTDPAPATTFAHLDATTVLSR.

## References

(1) Shekar, K. C.; Li, L.; Dabkowski, E. R.; Xu, W.; Ribeiro, R. F.; Hecker, P. A.; Recchia, F. A.; Sadygov, R. G.; Willard, B.; Kasumov, T.; Stanley, W. C. Cardiac Mitochondrial Proteome Dynamics with Heavy Water Reveals Stable Rate of Mitochondrial Protein Synthesis in Heart Failure despite Decline in Mitochondrial Oxidative Capacity. J. Mol. Cell. Cardiol. 2014, 75, 88–97. 10.1016/j.yjmcc.2014.06.014.

(2) Lau, E.; Cao, Q.; Ng, D. C. M.; Bleakley, B. J.; Dincer, T. U.; Bot, B. M.; Wang, D.; Liem, D. A.; Lam, M. P. Y.; Ge, J.; Ping, P. A Large Dataset of Protein Dynamics in the Mammalian Heart Proteome. Sci. Data 2016, 3, 160015. 10.1038/sdata.2016.15.

(3) Kim, T.-Y.; Wang, D.; Kim, A. K.; Lau, E.; Lin, A. J.; Liem, D. A.; Zhang, J.; Zong, N. C.; Lam, M. P. Y.; Ping, P. Metabolic Labeling Reveals Proteome Dynamics of Mouse Mitochondria. Mol. Cell. Proteomics MCP 2012, 11 (12), 1586–1594. 10.1074/mcp.M112.021162.

(4) Price, J. C.; Khambatta, C. F.; Li, K. W.; Bruss, M. D.; Shankaran, M.; Dalidd, M.; Floreani, N. A.; Roberts, L. S.; Turner, S. M.; Holmes, W. E.; Hellerstein, M. K. The Effect of Long Term Calorie Restriction on in Vivo Hepatic Proteostatis: A Novel Combination of Dynamic and Quantitative Proteomics. Mol. Cell. Proteomics 2012, 11 (12), 1801–1814. 10.1074/mcp.M112.021204.

(5) Busch, R.; Kim, Y.; Neese, R.; Schadeserin, V.; Collins, M.; Awada, M.; Gardner, J.; Beysen, C.; Marino, M.; Misell, L. Measurement of Protein Turnover Rates by Heavy Water Labeling of Nonessential Amino Acids. Biochim. Biophys. Acta BBA - Gen. Subj. 2006, 1760 (5), 730–744. 10.1016/j.bbagen.2005.12.023.

(6) Hammond, D. E.; Simpson, D. M.; Franco, C.; Wright Muelas, M.; Waters, J.; Ludwig, R. W.; Prescott, M. C.; Hurst, J. L.; Beynon, R. J.; Lau, E. Harmonizing Labeling and Analytical Strategies to Obtain Protein Turnover Rates in Intact Adult Animals. Mol. Cell. Proteomics 2022, 21 (7), 100252. 10.1016/j.mcpro.2022.100252.

(7) Miller, B. F.; Reid, J. J.; Price, J. C.; Lin, H.-J. L.; Atherton, P. J.; Smith, K. CORP: The Use of Deuterated Water for the Measurement of Protein Synthesis. J. Appl. Physiol. Bethesda Md 1985 2020, 128 (5), 1163–1176. 10.1152/japplphysiol.00855.2019.

(8) Lam, M. P. Y.; Wang, D.; Lau, E.; Liem, D. A.; Kim, A. K.; Ng, D. C. M.; Liang, X.; Bleakley, B. J.; Liu, C.; Tabaraki, J. D.; Cadeiras, M.; Wang, Y.; Deng, M. C.; Ping, P. Protein Kinetic Signatures of the Remodeling Heart Following Isoproterenol Stimulation. J. Clin. Invest. 2014, 124 (4), 1734–1744. 10.1172/JCI73787.

(9) Price, J. C.; Holmes, W. E.; Li, K. W.; Floreani, N. A.; Neese, R. A.; Turner, S. M.; Hellerstein, M. K. Measurement of Human Plasma Proteome Dynamics with (2)H(2)O and Liquid Chromatography Tandem Mass Spectrometry. Anal. Biochem. 2012, 420 (1), 73–83. 10.1016/j.ab.2011.09.007.

(10) Deberneh, H. M.; Abdelrahman, D. R.; Verma, S. K.; Linares, J. J.; Murton, A. J.; Russell, W. K.; Kuyumcu-Martinez, M. N.; Miller, B. F.; Sadygov, R. G. A Large-Scale LC-MS Dataset of Murine Liver Proteome from Time Course of Heavy Water Metabolic Labeling. Sci. Data 2023, 10 (1), 635. 10.1038/s41597-023-02537-w.

(11) Lau, E.; Cao, Q.; Lam, M. P. Y.; Wang, J.; Ng, D. C. M.; Bleakley, B. J.; Lee, J. M.; Liem, D. A.; Wang, D.; Hermjakob, H.; Ping, P. Integrated Omics Dissection of Proteome Dynamics during Cardiac Remodeling. Nat. Commun. 2018, 9 (1), 120. 10.1038/s41467-017-02467-3.

(12) Shankaran, M.; King, C. L.; Angel, T. E.; Holmes, W. E.; Li, K. W.; Colangelo, M.; Price, J. C.; Turner, S. M.; Bell, C.; Hamilton, K. L.; Miller, B. F.; Hellerstein, M. K. Circulating Protein Synthesis Rates Reveal Skeletal Muscle Proteome Dynamics. J. Clin. Invest. 2015, 126 (1), 288–302. 10.1172/JCI79639.

(13) Fanara, P.; Wong, P.-Y. A.; Husted, K. H.; Liu, S.; Liu, V. M.; Kohlstaedt, L. A.; Riiff, T.; Protasio, J. C.; Boban, D.; Killion, S.; Killian, M.; Epling, L.; Sinclair, E.; Peterson, J.; Price, R. W.; Cabin, D. E.; Nussbaum, R. L.; Brühmann, J.; Brandt, R.; Christine, C. W.; Aminoff, M. J.; Hellerstein, M. K. Cerebrospinal Fluid–Based Kinetic Biomarkers of Axonal Transport in Monitoring Neurodegeneration. J. Clin. Invest. 2012, 122 (9), 3159–3169. 10.1172/JCI64575.

(14) Deberneh, H. M.; Abdelrahman, D. R.; Verma, S. K.; Linares, J. J.; Murton, A. J.; Russell, W. K.; Kuyumcu-Martinez, M. N.; Miller, B. F.; Sadygov, R. G. Quantifying Label Enrichment from Two Mass Isotopomers Increases Proteome Coverage for in Vivo Protein Turnover Using Heavy Water Metabolic Labeling. Commun. Chem. 2023, 6 (1), 72. 10.1038/s42004-023-00873-x.

(15) Łącki, M. K.; Valkenborg, D.; Startek, M. P. IsoSpec2: Ultrafast Fine Structure Calculator. Anal. Chem. 2020, 92 (14), 9472–9475. 10.1021/acs.analchem.0c00959.

(16) Loos, M.; Gerber, C.; Corona, F.; Hollender, J.; Singer, H. Accelerated Isotope Fine Structure Calculation Using Pruned Transition Trees. Anal. Chem. 2015, 87 (11), 5738–5744. 10.1021/acs.analchem.5b00941.

(17) Dostal, V.; Wood, S. D.; Thomas, C. T.; Han, Y.; Lau, E.; Lam, M. P. Y. Proteomic Signatures of Acute Oxidative Stress Response to Paraquat in the Mouse Heart. Sci. Rep. 2020, 10 (1), 18440. 10.1038/s41598-020-75505-8.

(18) Sadygov, R. G. Partial Isotope Profiles Are Sufficient for Protein Turnover Analysis Using Closed-Form Equations of Mass Isotopomer Dynamics. Anal. Chem. 2020, 92 (21), 14747–14753. 10.1021/acs.analchem.0c03343.

(19) The UniProt Consortium; Bateman, A.; Martin, M.-J.; Orchard, S.; Magrane, M.; Ahmad, S.; Alpi, E.; Bowler-Barnett, E. H.; Britto, R.; Bye-A-Jee, H.; Cukura, A.; Denny, P.; Dogan, T.; Ebenezer, T.; Fan, J.; Garmiri, P.; Da Costa Gonzales, L. J.; Hatton-Ellis, E.; Hussein, A.; Ignatchenko, A.; Insana, G.; Ishtiaq, R.; Joshi, V.; Jyothi, D.; Kandasaamy, S.; Lock, A.; Luciani, A.; Lugaric, M.; Luo, J.; Lussi, Y.; MacDougall, A.; Madeira, F.; Mahmoudy, M.; Mishra, A.; Moulang, K.; Nightingale, A.; Pundir, S.; Qi, G.; Raj, S.; Raposo, P.; Rice, D. L.; Saidi, R.; Santos, R.; Speretta, E.; Stephenson, J.; Totoo, P.; Turner, E.; Tyagi, N.; Vasudev, P.; Warner, K.; Watkins, X.; Zaru, R.; Zellner, H.; Bridge, A. J.; Aimo, L.; Argoud-Puy, G.; Auchincloss, A. H.; Axelsen, K. B.; Bansal, P.; Baratin, D.; Batista Neto, T. M.; Blatter, M.-C.; Bolleman, J. T.; Boutet, E.; Breuza, L.; Gil, B. C.; Casals-Casas, C.; Echioukh, K. C.; Coudert, E.; Cuche, B.; De Castro, E.; Estreicher, A.; Famiglietti, M. L.; Feuermann, M.; Gasteiger, E.; Gaudet, P.; Gehant, S.; Gerritsen, V.; Gos, A.; Gruaz, N.; Hulo, C.; Hyka-Nouspikel, N.; Jungo, F.; Kerhornou, A.; Le Mercier, P.; Lieberherr, D.; Masson, P.; Morgat, A.; Muthukrishnan, V.; Paesano, S.; Pedruzzi, I.; Pilbout, S.; Pourcel, L.; Poux, S.; Pozzato, M.; Pruess, M.; Redaschi, N.; Rivoire, C.; Sigrist, C. J. A.; Sonesson, K.; Sundaram, S.; Wu, C. H.; Arighi, C. N.; Arminski, L.; Chen, C.; Chen, Y.; Huang, H.; Laiho, K.; McGarvey, P.; Natale, D. A.; Ross, K.; Vinayaka, C. R.; Wang, Q.; Wang, Y.; Zhang, J. UniProt: The Universal Protein Knowledgebase in 2023. Nucleic Acids Res. 2023, 51 (D1), D523–D531. 10.1093/nar/gkac1052.

(20) da Veiga Leprevost, F.; Haynes, S. E.; Avtonomov, D. M.; Chang, H.-Y.; Shanmugam, A. K.; Mellacheruvu, D.; Kong, A. T.; Nesvizhskii, A. I. Philosopher: A Versatile Toolkit for Shotgun Proteomics Data Analysis. Nat. Methods 2020, 17 (9), 869–870. 10.1038/s41592-020-0912-y.

(21) Eng, J. K.; Hoopmann, M. R.; Jahan, T. A.; Egertson, J. D.; Noble, W. S.; MacCoss, M. J. A Deeper Look into Comet--Implementation and Features. J. Am. Soc. Mass Spectrom. 2015, 26 (11), 1865–1874. 10.1007/s13361-015-1179-x.

(22) The, M.; MacCoss, M. J.; Noble, W. S.; Käll, L. Fast and Accurate Protein False Discovery Rates on Large-Scale Proteomics Data Sets with Percolator 3.0. J. Am. Soc. Mass Spectrom. 2016, 27 (11), 1719–1727. 10.1007/s13361-016-1460-7.

(23) Guan, S.; Price, J. C.; Ghaemmaghami, S.; Prusiner, S. B.; Burlingame, A. L. Compartment Modeling for Mammalian Protein Turnover Studies by Stable Isotope Metabolic Labeling. Anal. Chem. 2012, 84 (9), 4014–4021. 10.1021/ac203330z.

(24) Hellerstein, M. K.; Neese, R. A. Mass Isotopomer Distribution Analysis at Eight Years: Theoretical, Analytic, and Experimental Considerations. Am. J. Physiol.-Endocrinol. Metab. 1999, 276 (6), E1146–E1170. 10.1152/ajpendo.1999.276.6.E1146.

